# Accelerated Biochemical Kinetic Model Fitting via the Asynchronous, Generalized Island Method

**DOI:** 10.1101/660522

**Authors:** J Kyle Medley, Shaik Asifullah, Joseph Hellerstein, Herbert M Sauro

## Abstract

Mechanistic kinetic models of biological pathways are an important tool for understanding biological systems. Constructing kinetic models requires fitting the parameters to experimental data. However, parameter fitting on these models is a non–convex, non–linear optimization problem. Many algorithms have been proposed to addressing optimization for parameter fitting including globally convergent, population–based algorithms. The computational complexity of the this optimization for even modest models means that parallelization is essential. Past approaches to parameter optimization have focused on parallelizing a particular algorithm. However, this requires re–implementing the algorithm usinga distributed computing framework, which requires a significant investment of time and effort. There are two major drawbacks of this approach: First, the choice of best algorithm may depend on the model. Given the large variety of optimization algorithms available, it is difficult to re–implement every potentially useful algorithm. Second, when new advances are made in a given optimization algorithm, the parallel implementation must be updated to take advantage of these advantages. Thus, there is a continual burden placed on the parallel implementation. The drawbacks of re–implementing algorithms lead us to a different approach to parallelizing parameter optimization. Instead of parallelizing the algorithms themselves, we run many instances of the algorithm on single cores. This provides great flexibility as to the choice of algorithms by allowing us to reuse previous implementations. Also, it does not require the creation and maintenance of parallel versions of optimization algorithms. This approach is known as the island method. To our knowledge, the utility of the island method for parameter fitting in systems biology has not been previously demonstrated. For the parameter fitting problem, we allow islands to exchange information about their “best” solutions so that all islands leverage the discoveries of the few. This turns out to be avery effective in practice, leading to super–linear speedups. That is, if a single processor finds the optimal value of parameters in time *t*, then *N* processors exchanging information in this way find the optimal value much faster than *t/N*. We show that the island method is able to consistently provide good speedups for these problems. We also benchmark the island method against a variety of large, challenging kinetic models and show that it is able to consistently improve the quality of fit in less time than a single–threaded implementation.Our software is available at https://github.com/sys-bio/sabaody under a Apache 2.0 license.

---

Models in systems biology are based on varying degrees of granularity. Increased detail often comes at the expense of decreased scalability. For example, constraint–based models exhibit excellent scalability and are often used as a formalism for genome–scale reconstructions. This scalability is due in part to the fact that optimization of constraint–based models is ammenable to efficient solvers from linear programming (1). Kinetic models, however, are less scalable due to a larger number of parameters, more expensive objective function evaluations, and the presence of many local minima. However, kinetic models are necessary to study the mechanisms behind transient biological processes such as signaling cascades and circadian rhythms. Thus, there is an impetus to improve the optimization of kinetic models.

One class of algorithms that have been designed to solve non–convex, non–linear optimization problems is biologically–inspired population–based algorithms. A population–based algorithm maintains a pool of candidate solutions and improves the pool as a whole over time. Such algorithms are usually designed to balance speed of convergence with diversity of the solution candidates.

This class includes population–based methods such as genetic algorithms (GAs) (2), differential evolution (3), particle–swarm optimization (PSO) (4), harmony search (5), the artificial bee colony algorithm (ABC) (6), and many others.

The parallel nature of population–based methods lends these algorithms to efficient parallelization (7). However, modern cloud–based computing is best suited for algorithms that can be efficiently parallelized across nodes on a network. From an implementation perspective, this creates unique challenges that are not present in multithreaded parallelization such as: how much data must be exchanged between nodes, how much synchronization is required, and how can the system be made fault–tolerant? These issues can have a major effect on the scalability of the algorithm.

One approach to parallelizing population–based algorithms is to simply distribute the population update step across different nodes (8, 9). For example, in differential evolution, new candidate decision vectors can be evaluated on different nodes and later combined to form the complete population for a given iteration. This manner of parallelization offers a potentially linear performance increase, and for this reason it is often used in *multithreaded* parallelization. However, we discuss later that certain factors limit the scalability of using this same approach for distributed algorithms.

An alternative approach is the so–called generalized island method (8, 10–12) (also referred to as the island model; we avoid this terminology here because we reserve the word “model” for the biochemical models we are optimizing). The island method is a meta–algorithm that parallelizes optimization by running different copies of an optimization algorithm (or different algorithms) on different nodes in the network. This is based on the biologically–inspired principle of *punctuated equilibria* (13), which states that isolated, independently–evolving populations tend to quickly reach equilibrium, and that once genetic equilibrium is reached there is little further genetic drift, leading to stagnation.

In the island method, islands running on different nodes periodically exchange solutions, thereby preventing stagnation and leading to better solutions. If the migration is based on the best individuals in each island, the method also exhibits accelerated convergence compared to a single island working alone. Additionally, the island method allows reusing previously existing algorithm implementations. The PaGMO2 library (11, 14), which we use in this work, provides implementations for 14 global and 16 local fitting algorithms, which was possible in part because of pre-existing implementations of the algorithms.

The island method has been applied to various diverse optimization problems such as nuclear power plant feedwater monitoring (15), vehicle routing (16), and trajectory planning for spacecraft (8, 11). The island method has also been applied to large–scale kinetic systems biology models (17–20). In this article, we apply the island method to a set of very large, challenging dynamical models from systems biology (21) in order to quantify the performance of different algorithms and the performance gains due to parallelization using the island model. We show that the distributed island method is able to accelerate parameter fitting and obtain better solutions for all models tested. As expected, the island method scales sub–linearly, but nevertheless provides a substantial improvement to performance and quality of fit. Our approach is completely asynchronous and uses very little network bandwidth, making it in principle an ideal candidate for scalling to massive numbers of nodes, even if the maximum bandwidth of the network infrastructure is low. We provide a reusable version of our software at https://github.com/distrib-dyn-modeling/sabaody. This software can be easily installed as a docker container on cloud services such as Amazon Web Services (AWS) and Google Compute Engine.

## Approach

Our main objective in this study was to test whether the island method can improve the convergence time and solution quality for large, challenging dynamical models, and to quantify the performance of different algorithms and algorithm combinations. In order to perform these tests, we constructed a series of three benchmarks. In the first benchmark, we reduced the combinatorial complexity of the different configuration options for the island method. This allowed us to run the remaining benchmarks in a realistic timeframe. In the second benchmark, we tested the scaling and convergence performance of the island method on our most challenging problems. Finally, in the third benchmark, we looked at the performance of different algorithms and combinations. Partitioning our benchmarks in this way allowed us to test our main objectives in turn while keeping the number of different configurations within feasible limits.

The island method is highly configurable. In addition to specifying potentially different fitting algorithms for each island, the migration routes between the islands can be connected in any arbitrary topology. Although our system allows for fully customizable user–specified migration topologies, we provide a set of 15 default topologies that can be generated automatically for any island number. These topologies are listed in Table 2. Since PaGMO2 contains 14 global and 16 local optimization algorithms, there are 30 · 15 = 450 algorithm / topology combinations to choose from for a given number of islands, not including algorithm combinations. Additionally, the island method can be used with different migration policies. In this work, we use a policy that selects the best *M* individuals from a source island, and replaces the worst *M* individuals in the destination island if the replacement candidate has a better (lower) score. Another strategy is random selection and replacement, which can lead to more genetic diversity but has a less pronounced acceleration effect.

Faced with this large combination of options, we created a systematic test to eliminate configurations that were unlikely to perform well. We constructed a set of 1120 benchmarks using different algorithm and topology combinations based on five analytic objective functions plotted in two– dimensions in Figure 7. These analytic functions have a much lower computational cost than our biochemical benchmark models, and are often used to evaluate newly developed nonlinear optimization algorithms. We refer to this set of tests as the elimination benchmark.

Table 3 shows that algorithm choice is more strongly correlated with performance than topology in our tests. Therefore, we focused on the effect of algorithm choice for the remaining benchmarks. We first ran benchmarks of problems B1 and B3 from the BioPreDyn suite (21) on the same topology / algorithm combination while varying the number of islands. These problems are expensive, but also allow us to critically test the convergence properties and performance of the island method. Next, we ran different algorithm combinations on problems B2 and B4 from BioPreDyn. Our purpose this time was to test for trends in algorithm performance, and to investigate whether a “synergistic” effect could be observed from combinations of different algorithms.

## Methods

### Elimination Benchmark

Ten–dimensional versions of the functions in Figure 7 were run for three replicates on the combinations of topologies and algorithms shown here. In some cases, algorithm combinations were used, such as `de+sade`. In such cases, the algorithms were alternated along the topology structure if linear or ring–shaped, or arbitrarily distributed otherwise (e.g. hypercube and random graph–based models). However, once assigned, the algorithm positions were not changed for the duration of all benchmarks. Results were grouped by topology and algorithm/combination respectively and independently ranked for each of the five problems according to the total number of rounds required for convergence. A lower score implies that the benchmark finished in fewer rounds. This benchmark consisted of up to 2000 rounds of migration, each interspersed with 1000 iterations of the local algorithm on each node. The fitting problem was terminated when the MSE of any decision vector dropped below a cutoff value of 0.01 with respect to the best known solution. Migration was fully asynchronous. The source data for this ranking is available in the supplementary information at TODO: upload data.

### Speedup Benchmark

In order to quantify the speed improvement of the island method, we benchmarked the B3 model for various numbers of islands using a constant *total population*. For the single–island case, the population of the island was 4800 individuals. In the case of two islands, the population of each island was 2400 individuals, and so on. This is a fair method of measuring speedup because a property of population–based algorithms is that the population must be suitably large in order to sufficiently explore the parameter space. This threshold is problem–dependent, and is usually scaled with the number of parameters. In this work, the total population was chosen based on the minimum value that tended to avoid getting stuck in local minima. Figure **??** shows the results of the speedup benchmark.

### Large Model Scalability Benchmark

In order to quantify the performance improvement of the island method, we benchmarked problems B1 and B3 from the BioPreDyn suite against different numbers of parallel islands. B1 is a genome scale *S. cerevisiae* model whereas B3 is a model of central carbon metabolism and transcription in *E. coli*. Both of these are large and challenging models, with the original authors reporting ≈ 1–week fitting times for both (21). Figure 4 shows that reasonable fits can be obtained in a day for B3 on our 16–core cluster.

**Fig. 1.**
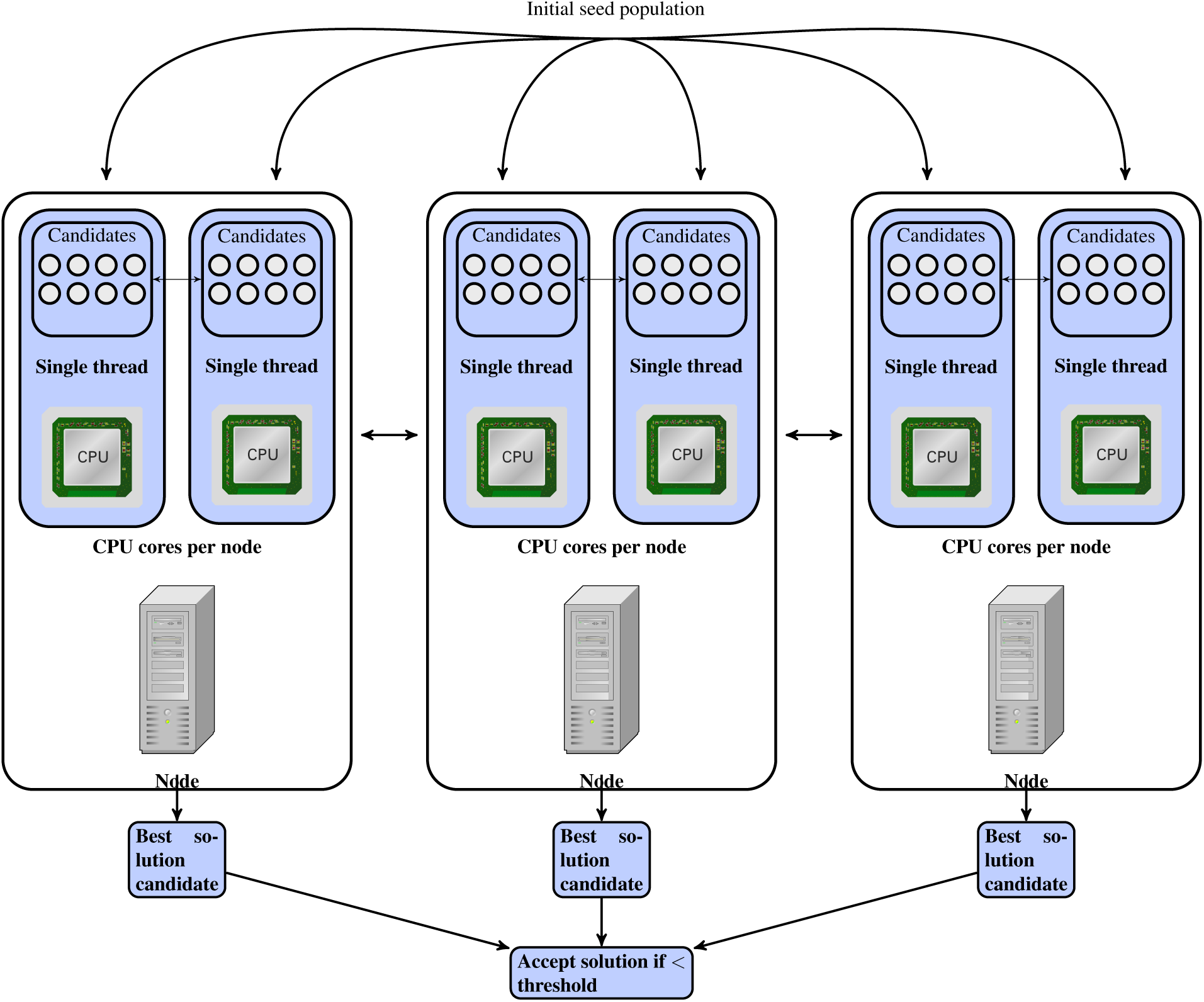
A visual depiction of the island method. A cluster containing a number of nodes (machines) and CPUs (one per processor core on each machine) is initialized randomly with a seed population. Each CPU maintains a population of solution vectors corresponding to one island. At the end of *k* iterations of the local algorithm (which can be different for each CPU), the best solutions obtained so far are exchanged among CPUs on the same or different machines.

**Fig. 2.**
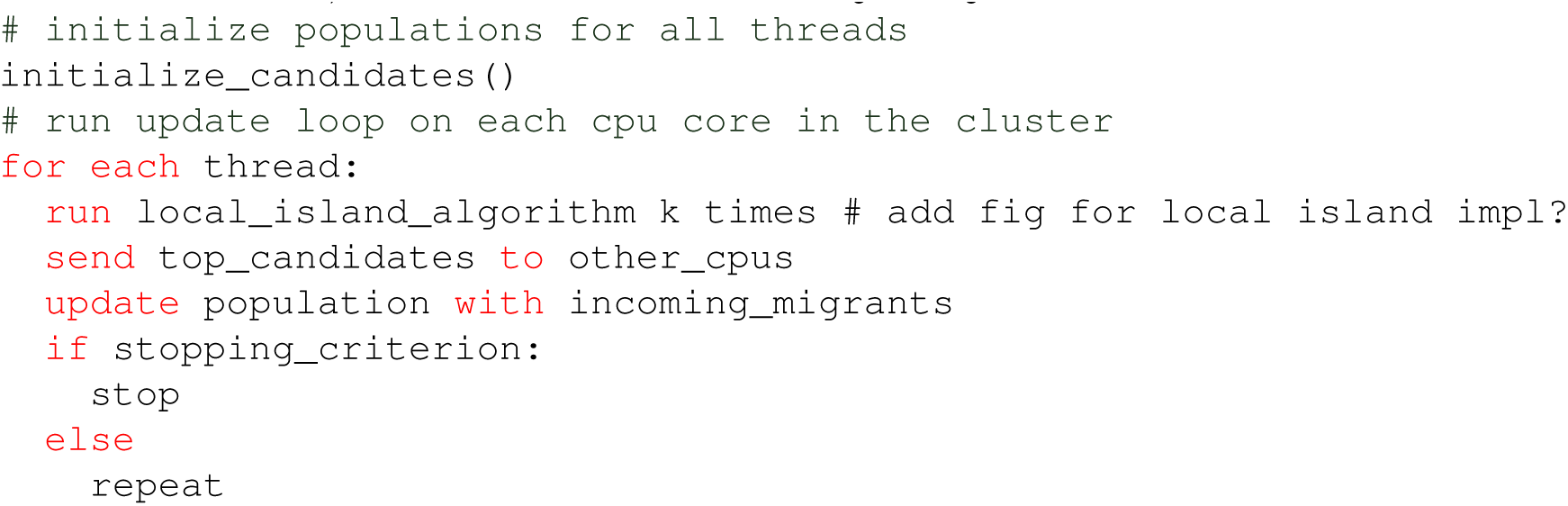
Distributed island method pseudocode. The island method runs locally on a number of threads, which can be on different CPU cores or different machines (the total number of threads should not exceed the total number of CPU cores in the cluster). In every update loop, the island method first runs the local algorithm (e.g. DE, ABC, etc.) for *k* iterations on each thread separately. Next, each island *i* sends its top *l* solutions to neighboring islands and receives up to *l × m* solutions from neighboring islands, where *m* is the connectivity of island *i*. This process repeats until a stopping criterion is reached (the objective function reaches a threshold value or the maximum number of iterations is exhausted).

**Fig. 3.**
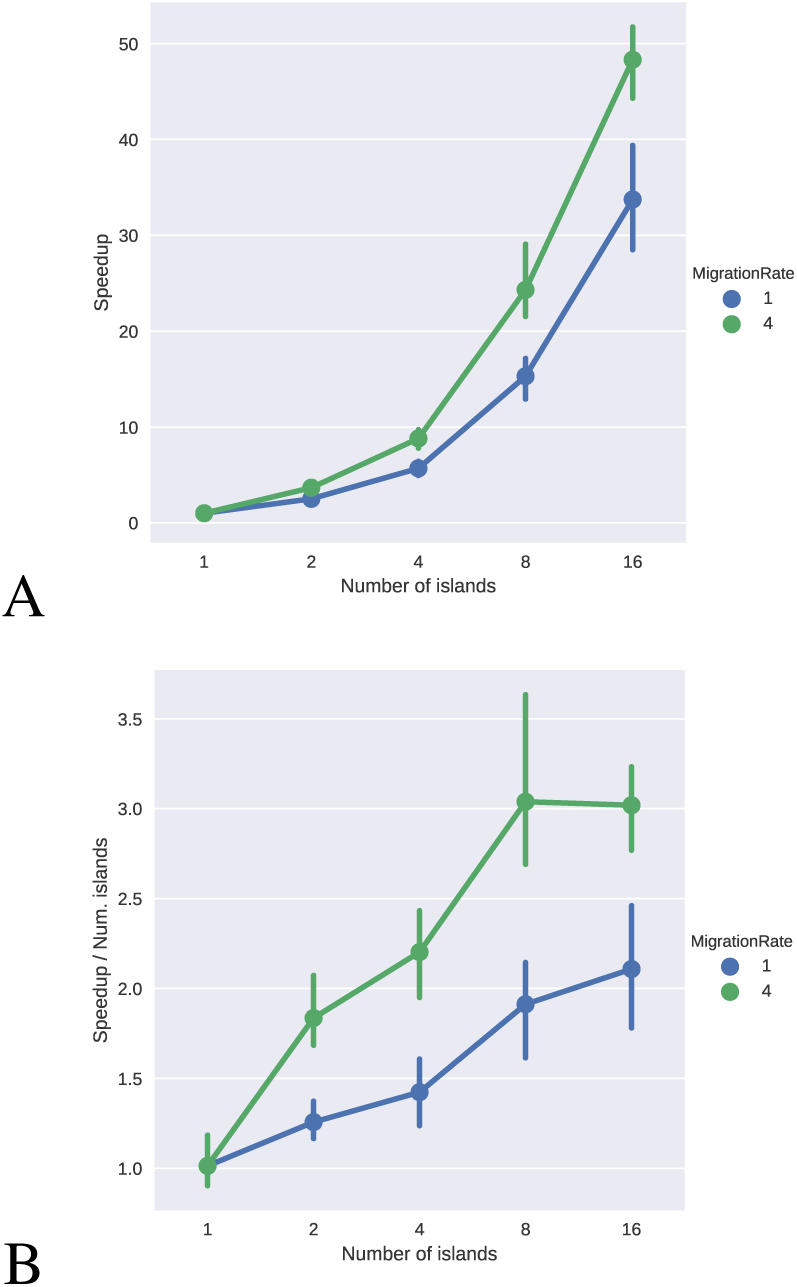
Speedup for the island method. Benchmarks were performed against the B3 problem using a constant total population (4800) for all islands (i.e. the sum of the populations of individual islands is constant) for three replicates. To enable running single–island variants in a reasonable timeframe (≈ 60 hrs for the single–island case), we reduced the box–constrained range for all log_10_ parameter values from ±1 to ±0.1. The convergence and termination criteria were as in Figure 4B. We expected the island method to exhibit sub–linear scaling behavior, but surprisingly, the opposite was true. A 16–island benchmark (using 16 cores) converged more than 40*×* faster than the single–core / single–island case **(A)**. This effect was quantified for two different migration rates (1/island/round and 4/island/round, resp.). All other benchmarks reported here use a migration rate of 4/island/round. To quantify this scaling, we plotted the same datapoints divided by the respective number of islands **(B)**. In this plot, linear scaling would be plotted as a horizontal line with a value of 1. However, the island method appears to operate at 3*×* this factor for large numbers of islands. A possible explanation for this result is provided in the Discussion Section.

**Fig. 4.**
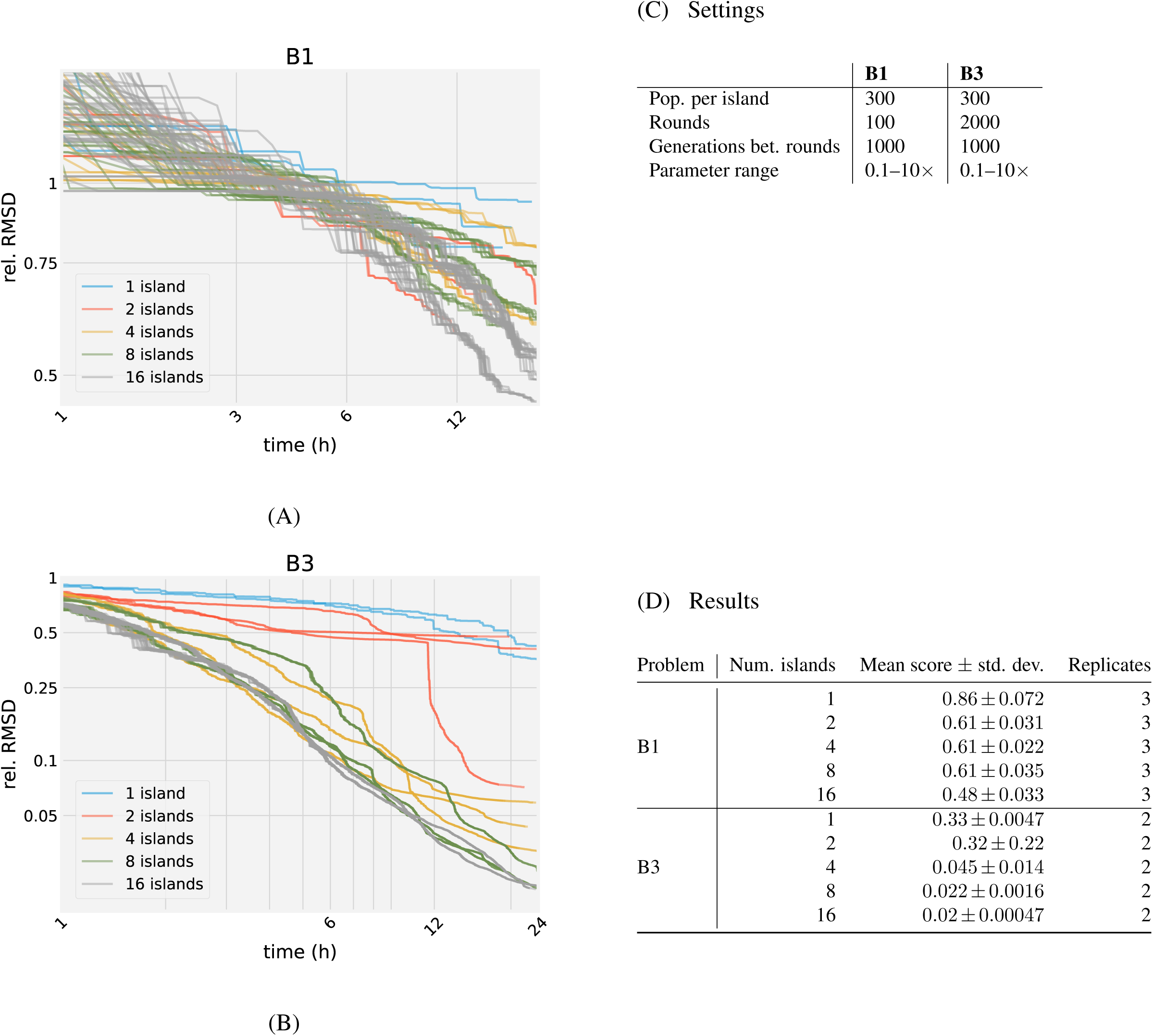
Convergence curves for problems B1 (A) and B3 (B) from the BioPreDyn suite (21) for various numbers of islands. Each curve plots the best *champion fitness* (i.e. the best solution up to the current time) per island over time. The settings for each benchmark are shown (C). For a given number of rounds, table (D) shows that increasing the island size yields an improvement in fitted parameters. As with BioPreDyn, we constrained the values of all fitted parameters to 0.1–10*x* the nominal value. The fitness value in (A) and (B) is computed from the root–mean–square deviation for each state variable normalized by the state variable’s average (in B1, the values are also rescaled to balance error contributions). Not all islands finish at the same time due to our use of variable–step integration in SBML simulation.

### Algorithm Benchmark

Table 3 suggests that combining local and global fitting algorithms can lead to better performance than either type of algorithm individually. For example, `de+nelder mead`, `de+praxis`, `de+sade`, and `de+de1220` are all ranked better than `de` alone. To test whether this trend would hold for more realistic fitting problems, we benchmarked problems B2 and B4 from the BioPre-Dyn suite against different algorithms and algorithm combinations, and quantified the results in Table 1. Figures 5 and 6 show the value of the current champion (the individual with the best fitness) in the population over time.

**Fig. 5.**
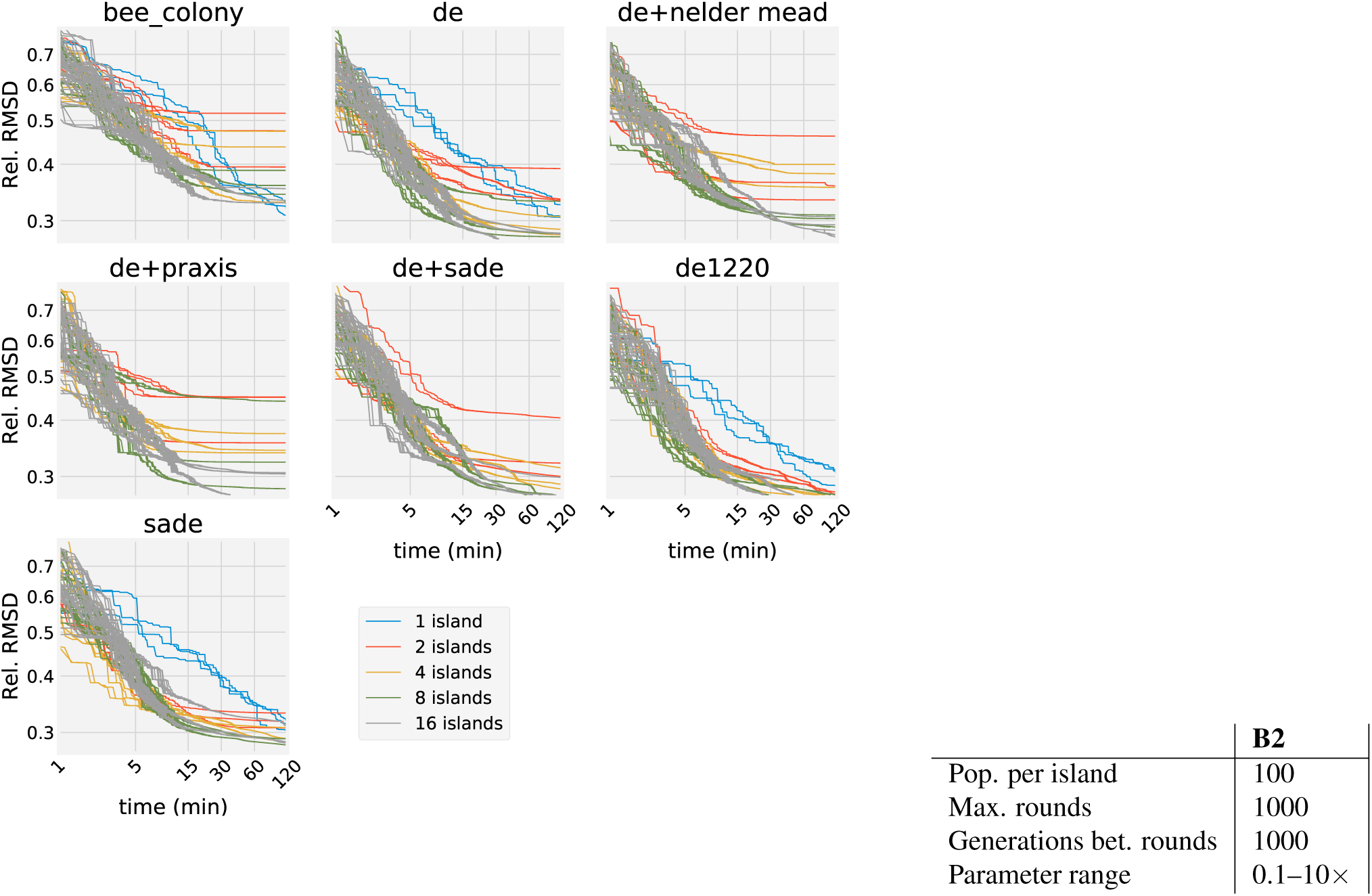
Convergence curves for problem B2 from the BioPreDyn suite. To test the effect of different algorithms and combinations of algorithms on the convergence rate, we tested the top performing algorithms and combinations determined from the elimination benchmark. Each trace represents the champion value (the individual with the best fitness) for a single island running on 1 CPU core. Single island variants are elided from combination benchmarks containing two or more algorithms.

**Fig. 6.**
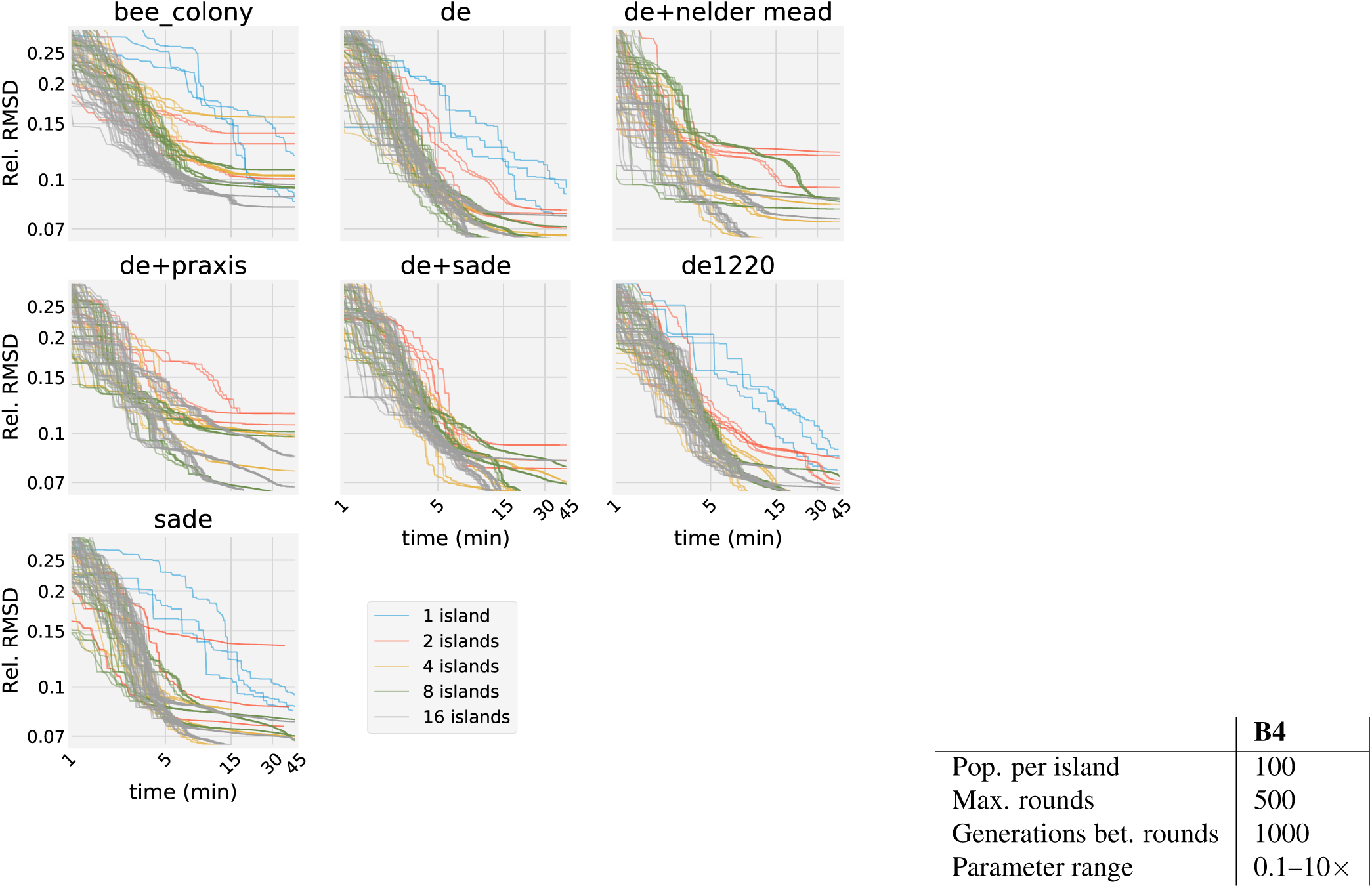
Convergence curves for problem B4 from the BioPreDyn suite. As with Figure 5, we plotted the champion values over time for each benchmark configuration.

## Results & Discussion

### Scaling Behavior

Figure 3 shows that the island method exhibits superlinear scaling. Parallel implementations of algorithms have been shown to exhibit this property due to more effective use of limited CPU cache resources (), but this would not account for the large ≈ 3× superlinearity factor observed here. Instead, the effect observed here is most likely due to the influence of the migration operator on the solution population. It is well–known that genetic and swarm– based algorithms can be tuned to trade population diversity for convergence rate by adjusting their respective algorithmic tuning parameters. The migration rate can be thought of as another tuning parameter. Figure 3 shows a strong dependence of the superlinear speedup on migration rate. In our implementation, islands always select their best solutions (in terms of objective value) as migrants. Thus, the population fraction of these solutions tends to increase over time and cause clustering of the population around the best minimum found so far. However, too much of this effect causes the island method to get stuck in local minima. Thus, the migration rate must be carefully tuned to balance these two effects. In our benchmarks, a very small rate of migration (4 per island per round) was sufficient to generate a considerable speedup. This can be understood in light of superlinear effects seen in other types of problems, such as depth–first search (22). Since some instances of the search algorithm will discover the optimal solution more quickly due simply to better initial conditions, algorithms which are sensitive to initial conditions (such as depth–first search and parameter optimization) can be expected to benefit from coordination among parallell instances of the algorithm. This suggests that the main benefit of migration is to allow certain islands to “share” good candidate solutions that are discovered simply because the island happened to be initialized with good initial conditions.

The fact that the island method is able to achieve good speedups using a very small migration rate is another desirable property. Whereas parallelization of the main loop may be an effective way to speed up population–based algorithms (9), it also requires more network bandwidth. For example, differential evolution forms new decision vectors from a linear combination of three individuals from the prior iteration. For a population of size *M* with *N* nodes, each node will evaluate *M/N* decision vectors and require 3*M/N* prior decision vectors to be sent over the network interface, which imposes bandwidth requirements for the network infrastructure. In our benchmarks, we used up to 2 million total iterations (2000 rounds of 1000 iterations) across a total population size of 16 · 300 = 4800. Assuming a decision vector of length 100, double–precision, this equates to 2.5 TB (terabytes) of information that must be sent over the network. In comparison, the island method, using a migration rate of 4 individuals per island per round, would require only 4 · 16 · 2000 · 100 · 64 = 100*MB*.

### Comparison with Distributed Algorithms

Most population–based algorithms can be implemented in a distributed fashion, by spreading the update step across *N* cores or nodes. This requires re-implementing the algorithm using a distributed computing framework, but can provide potentially linear speedups. For example, when using 16 total CPUs (the sum of all CPUs in the cluster), a distributed algorithm would be expected to be ≈ 16× faster. In contrast, the speedup provided by the island method is dependent on the frequency of migration between nodes. However, increasing the migration frequency excessively causes a drop in population diversity due to the fact that the selection policy always chooses the best candidate solution in a population, hence enriching the population fraction of this candidate over time. Thus, in general, the island method requires careful tuning to achieve linear speedups, whereas a distributed implementation of a single algorithm would be expected to exhibit these speedups automatically. This may be seen as a disadvantage of the island model. However, this is somewhat offset by the fact that, in all our benchmarks, we were able to achieve good performance using a constant migration rate of 4 individuals per round. Thus, just by migrating a small fraction of individuals in comparison to the population size per island (which was 100–300 in all benchmaarks), we were able to attain good speedups without needing to tune the migration rate. Furthermore, this illustrates the fact that the island method requires significantly less bandwidth than a distributed fitting algorithm, which is a major consideration for scalability.

For example, differential evolution forms new decision vectors from a linear combination of three individuals from the prior iteration. For a population of size *M* with *N* nodes, each node will evaluate *M/N* decision vectors and require 3*M/N* prior decision vectors to be sent over the network interface, which imposes bandwidth requirements for the network infrastructure. In our benchmarks, we used up to 2 million total iterations (2000 rounds of 1000 iterations) across a total population size of 16 · 300 = 4800. Assuming a decision vector of length 100, double–precision, this equates to 20 Tb (terabits) of information that must be sent over the network. In comparison, the island method using a migration rate of 4 individuals per island per round would require only 4 · 16 · 2000 · 100 · 64 = 820*Mb*. Whether or not these requirements pose a limtation depends on the networking infrastructure used.

More importantly, distributed optimization algorithms must be re-implemented using a distributed computing framework such as Spark (23) or Dask (24). In many cases, an independently–validated reference implementation of an algorithm is available in C/C++ for running locally on a single node, but not on distributed nodes. Thus, if a researcher wishes to evaluate a given fitting algorithm using distributed computing, the researcher must first re-implement the algorithm itself using distributed technologies. Since computing frameworks like Spark (23) or Dask (24) are evolving rapidly, it is not known whether a re-implementation will be rendered obsolete by technology changes, thereby re-creating the same dilemma again in the future.

### Hardware Specification

Our hardware used for this study consisted of two workstations with a 10–core Intel^®^ Xeon^®^ CPU E5–2660V3 at 2.6 GHz with 24 GB RAM and a 6– core Intel^®^ Xeon^®^ CPU E5–2620V2 at 2.1 GHz with 64 GB RAM.

## Conclusion

Model calibration is often a major bottleneck in constructing dynamical models. The ability to accelerate this process would allow modelers more freedom to experiment with different model variants, and shorten the overall length of model development. We have shown here that the asynchronous, distributed island method yields accelerated convergence and better quality of fit for very large, challenging optimization problems in systems biology.

We also tested the B2 and B4 algorithms against a range of algorithm types. The results shown here indicate that there is little variability in algorithm performance for different problems, and that combinations of different algorithm types (in a heterogeneous island configuration) does not significantly enhance performance. Previously, Villaverde *et al.* (25) reported a sequential / temporal approach to implementing hybrid algorithms (rather than a migration–based approach, as in the case of the island method). By using adjoint–based sensitivities for computing the local gradient combined with a global scatter search metaheuristic, Villaverde *et al.* obtained significant performance improvements on large models using a hybrid approach. In our work, we do not observe performance improvements when combining local and global algorithms. This discrepancy may be due to (1) the nature of the algorithms tested (we do not consider adjoint–based gradient methods, which were an important component of the hybrid strategy used by Villaverde *et al.*), (2) the sequential vs topological partitioning strategy used by the different approaches, or (3) the enrichment effect of the “best candidate” migration strategy used here, which exacerbates the effect of local minima. A thorough examination of the convergence properties of both approaches, using identical local and global algorithms in each case, might reveal interesting trends in the efficacy of different methods for hybrid optimization under different conditions, and may suggest general strategies for combining local and global algorithms.

A common criticism of biologically–inspired population– based methods is that such algorithms require careful tuning of algorithm parameters such as crossover rate (for genetic algorithms) and local versus global attraction (swarm–based methods). However, these results show that topology and algorithm choice only exhibit a minor role in determining performance, at least for the benchmarks tested here. This suggests that the island method is useful as a general parallelization scheme without the need to tune these hyperparameters. The top–performing combination of the *de1220* algorithm with the rim topology exhibits good general performance and scalability, and should be sufficient for most users. Users can scale this combination by setting the number of islands equal to the total number of CPU cores in the cluster.

## Acknowledgements

The authors would like to express their sincere gratitude to the Google and the National Resource for Network Biology for providing funding for this project through the Google Summer of Code program.

## Supplementary Information

**Table 1.** Results for problems B2 and B4.

| Problem | Algorithm | Num. islands | Mean score ± std. dev. | Hours | Rounds | Converged? | Replicates |
| --- | --- | --- | --- | --- | --- | --- | --- |
| B2 | bee colony | 1 | $0.3 \pm 0.0091$ | $3.1 \pm 0.11$ | $1000.0 \pm 0.0$ | No | 3 |
| | | 16 | $0.34 \pm 0.013$ | $5.4 \pm 1.8$ | $1000.0 \pm 0.0$ | No | 3 |
| | de | 1 | $0.31 \pm 0.011$ | $2.0 \pm 0.0042$ | $1000.0 \pm 0.0$ | No | 3 |
| | | 16 | $0.28 \pm 0.004$ | $2.0 \pm 1.2$ | $740.0 \pm 450.0$ | Yes | 3 |
| | de+nelder mead | 1 | $0.32 \pm 0.0075$ | $2.0 \pm 0.026$ | $1000.0 \pm 0.0$ | No | 3 |
| | | 16 | $0.29 \pm 0.016$ | $3.1 \pm 0.86$ | $930.0 \pm 130.0$ | Yes | 3 |
| | de+praxis | 1 | $0.32 \pm 0.011$ | $2.3 \pm 0.0031$ | $1000.0 \pm 0.0$ | No | 3 |
| | | 16 | $0.29 \pm 0.019$ | $2.0 \pm 1.2$ | $770.0 \pm 400.0$ | Yes | 3 |
| | de+sade | 1 | $0.32 \pm 0.0053$ | $2.2 \pm 0.013$ | $1000.0 \pm 0.0$ | No | 3 |
| | | 16 | $0.28 \pm 0.012$ | $1.9 \pm 0.88$ | $660.0 \pm 300.0$ | Yes | 3 |
| | de1220 | 1 | $0.3 \pm 0.012$ | $2.0 \pm 0.049$ | $1000.0 \pm 0.0$ | No | 3 |
| | | 16 | $0.28 \pm 0.012$ | $0.62 \pm 0.19$ | $200.0 \pm 120.0$ | Yes | 3 |
| | sade | 1 | $0.31 \pm 0.0068$ | $2.3 \pm 0.028$ | $1000.0 \pm 0.0$ | No | 3 |
| | | 16 | $0.28 \pm 0.0011$ | $3.3 \pm 1.0$ | $1000.0 \pm 0.0$ | No | 3 |
| B4 | bee colony | 1 | $0.08 \pm 0.0043$ | $1.8 \pm 0.71$ | $500.0 \pm 0.0$ | No | 3 |
| | | 2 | $0.12 \pm 0.02$ | $1.1 \pm 0.059$ | $500.0 \pm 0.0$ | No | 3 |
| | | 4 | $0.12 \pm 0.031$ | $1.1 \pm 0.007$ | $500.0 \pm 0.0$ | No | 3 |
| | | 8 | $0.098 \pm 0.008$ | $1.1 \pm 0.023$ | $500.0 \pm 0.0$ | No | 3 |
| | | 16 | $0.089 \pm 0.0073$ | $1.1 \pm 0.034$ | $500.0 \pm 0.0$ | No | 3 |
| | de | 1 | $0.086 \pm 0.0074$ | $0.76 \pm 0.027$ | $500.0 \pm 0.0$ | No | 3 |
| | | 2 | $0.076 \pm 0.0051$ | $0.84 \pm 0.067$ | $500.0 \pm 0.0$ | No | 3 |
| | | 4 | $0.07 \pm 0.006$ | $0.84 \pm 0.061$ | $500.0 \pm 0.0$ | No | 3 |
| | | 8 | $0.067 \pm 0.0034$ | $0.63 \pm 0.35$ | $380.0 \pm 200.0$ | Yes | 3 |
| | | 16 | $0.069 \pm 0.0069$ | $0.47 \pm 0.34$ | $290.0 \pm 200.0$ | Yes | 3 |
| | de+nelder mead | 1 | $0.089 \pm 0.0046$ | $0.78 \pm 0.035$ | $500.0 \pm 0.0$ | No | 3 |
| | | 2 | $0.11 \pm 0.015$ | $0.82 \pm 0.062$ | $500.0 \pm 0.0$ | No | 3 |
| | | 4 | $0.074 \pm 0.0091$ | $0.75 \pm 0.091$ | $490.0 \pm 15.0$ | Yes | 3 |
| | | 8 | $0.084 \pm 0.0034$ | $0.79 \pm 0.008$ | $500.0 \pm 0.0$ | No | 3 |
| | | 16 | $0.076 \pm 0.011$ | $0.58 \pm 0.37$ | $370.0 \pm 220.0$ | Yes | 3 |
| | de+praxis | 1 | $0.085 \pm 0.0031$ | $0.77 \pm 0.032$ | $500.0 \pm 0.0$ | No | 3 |
| | | 2 | $0.11 \pm 0.0053$ | $0.98 \pm 0.31$ | $500.0 \pm 0.0$ | No | 3 |
| | | 4 | $0.09 \pm 0.013$ | $0.82 \pm 0.045$ | $500.0 \pm 0.0$ | No | 3 |
| | | 8 | $0.088 \pm 0.02$ | $0.81 \pm 0.056$ | $500.0 \pm 3.0$ | No | 3 |
| | | 16 | $0.072 \pm 0.0098$ | $0.64 \pm 0.29$ | $410.0 \pm 160.0$ | Yes | 3 |
| | de+sade | 1 | $0.088 \pm 0.0028$ | $0.76 \pm 0.034$ | $500.0 \pm 0.0$ | No | 3 |
| | | 2 | $0.084 \pm 0.0074$ | $0.82 \pm 0.0099$ | $500.0 \pm 0.0$ | No | 3 |
| | | 4 | $0.066 \pm 0.0024$ | $0.51 \pm 0.27$ | $330.0 \pm 150.0$ | Yes | 3 |
| | | 8 | $0.071 \pm 0.007$ | $0.67 \pm 0.29$ | $410.0 \pm 150.0$ | Yes | 3 |
| | | 16 | $0.07 \pm 0.0098$ | $0.41 \pm 0.34$ | $260.0 \pm 210.0$ | Yes | 3 |
| | de1220 | 1 | $0.083 \pm 0.0064$ | $0.71 \pm 0.0091$ | $500.0 \pm 0.0$ | No | 3 |
| | | 2 | $0.074 \pm 0.0075$ | $0.8 \pm 0.024$ | $500.0 \pm 0.0$ | No | 3 |
| | | 4 | $0.07 \pm 0.0086$ | $0.21 \pm 0.11$ | $140.0 \pm 74.0$ | Yes | 3 |
| | | 8 | $0.067 \pm 0.0034$ | $0.47 \pm 0.28$ | $300.0 \pm 170.0$ | Yes | 3 |
| | | 16 | $0.066 \pm 0.00095$ | $0.67 \pm 0.31$ | $400.0 \pm 170.0$ | Yes | 3 |
| | sade | 1 | $0.088 \pm 0.0051$ | $0.78 \pm 0.085$ | $500.0 \pm 0.0$ | No | 3 |
| | | 2 | $0.099 \pm 0.032$ | $0.63 \pm 0.033$ | $500.0 \pm 0.0$ | No | 3 |
| | | 4 | $0.073 \pm 0.01$ | $0.42 \pm 0.3$ | $280.0 \pm 190.0$ | Yes | 3 |
| | | 8 | $0.072 \pm 0.006$ | $0.83 \pm 0.043$ | $500.0 \pm 0.0$ | No | 3 |
| | | 16 | $0.07 \pm 0.0067$ | $0.64 \pm 0.31$ | $390.0 \pm 190.0$ | Yes | 3 |

**Table 2.** Topology presets included with our implementation. These presets can be used to generate a topology for any number of islands (except the hypercube, which requires that the number of islands be a power of two).More detailed information, including topology illustrations, is available at https://sabaody.readthedocs.io/en/latest/topologies.html

| Topology |
| --- |
| One-way Ring |
| Bidirectional ring |
| Bidirectional chain |
| Lollipop |
| Rim |
| 1-2 Ring |
| 1-2-3 Ring |
| Fully Connected (Complete Graph) |
| Broadcast |
| Hypercube |
| Watts-Strogats |
| Erdős-Rényi |
| Barabási-Albert |
| Extended Barabási-Albert |
| Extended Ageing Barabási-Albert |

**Table 3.** Ranking of the elimination benchmark problems. The bottom row shows the mean difference between the highest and lowest ranking

| Topology | Range | Algorithm | Range |
| --- | --- | --- | --- |
| Hypercube | 1–9 | de1220 | 2–4 |
| Rim | 2–8 | de+nelder mead | 1–8 |
| Bidirectional ring | 4–7 | bee colony | 1–10 |
| Bidirectional chain | 1–11 | de+praxis | 2–9 |
| Barabási–Albert (26) | 1–13 | de+sade | 4–7 |
| 1–2 Ring | 3–12 | sade | 2–9 |
| Broadcast | 1–14 | de+de1220 | 4–9 |
| Fully Connected | 2–13 | de | 6–8 |
| One–way ring | 7–8 | de+pso | 7–10 |
| 1–2–3 Ring | 4–12 | de+nsga2 | 8–10 |
| Ageing Extended Barabási–Albert | 6–10 | pso | 11–14 |
| Erdős–Rényi (27) | 3–14 | ihs | 11–14 |
| Extended Barabási–Albert | 5–12 | xnes | 11–14 |
| Watts–Strogatz (28) | 5–14 | nsga2 | 12–14 |
| **Avg. diff.** | **8.00** | **Avg. diff.** | **3.81** |

**Fig. 7.**
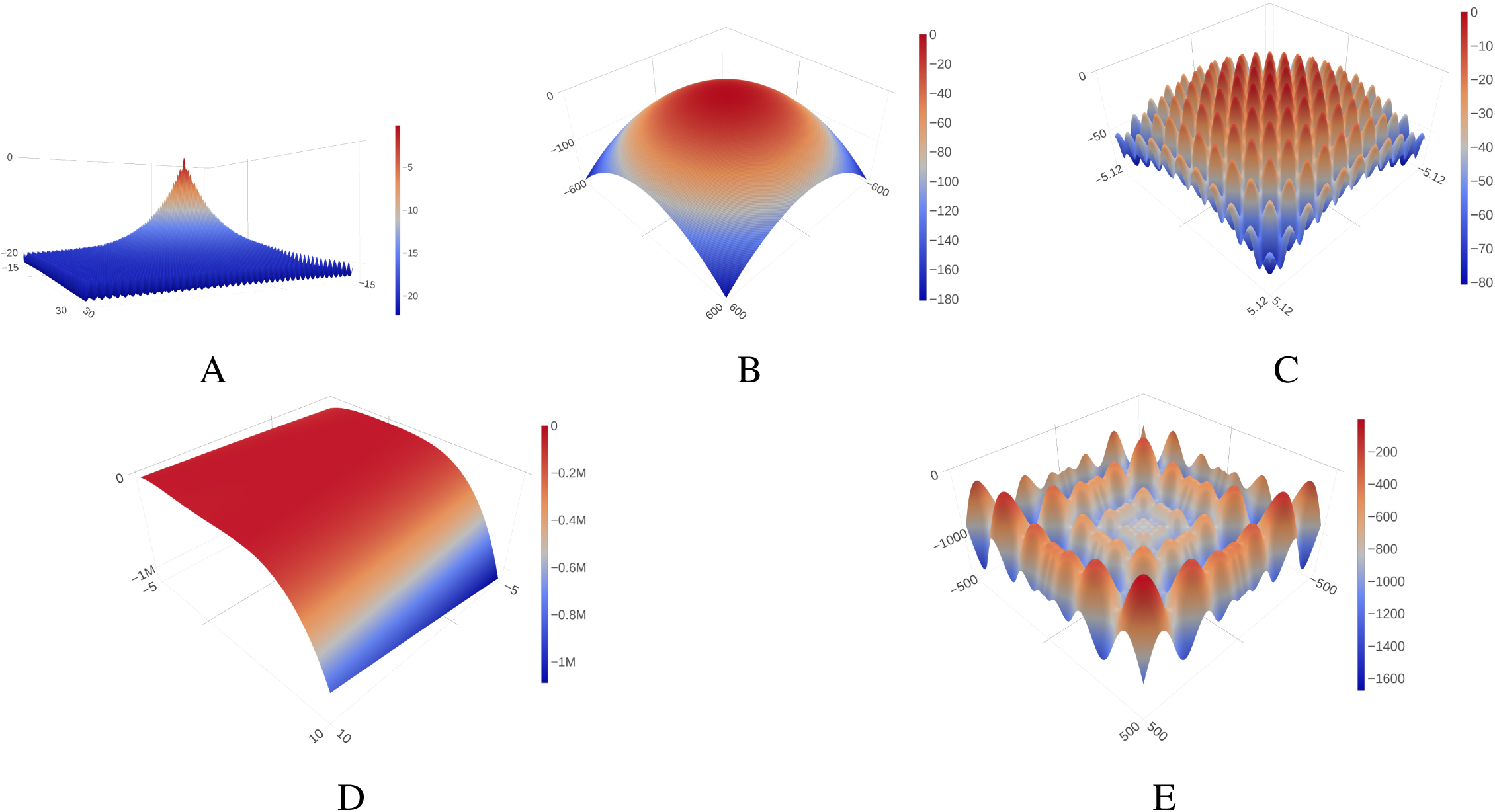
Two–dimensional plots of our five elimination benchmark functions. These functions are analytical, and hence allow for testing a much larger number of combinations than our biochemical model benchmarks. The functions are analytic closed–form expressions: Ackley (A) (29), Griewank (B) (30), Rastrigin (C) (31), Rosenbrock (D) (32), and Schwefel (E) (33) functions. The plots are inverted so that the minimum value of the function is at the highest elevation, which allows for better visualization.

